# Altered brain criticality in Schizophrenia: New insights from MEG

**DOI:** 10.1101/2021.11.09.467906

**Authors:** Golnoush Alamian, Tarek Lajnef, Annalisa Pascarella, Jean-Marc Lina, Laura Knight, James Walters, Krish D Singh, Karim Jerbi

## Abstract

Schizophrenia has a complex etiology and symptomatology that is difficult to untangle. After decades of research, important advancements towards a central biomarker are still lacking. One of the missing pieces is a better understanding of how non-linear neural dynamics are altered in this patient population. In this study, the resting-state neuromagnetic signals of schizophrenia patients and healthy controls were analyzed in the framework of criticality. When biological systems like the brain are in a state of criticality, they are thought to be functioning at maximum efficiency (e.g., optimal communication and storage of information) and with maximum adaptability to incoming information. Here, we assessed the self-similarity and multifractality of resting-state brain signals recorded with magnetoencephalography in patients with schizophrenia patients and in matched controls. Our analysis showed a clear ascending, rostral to caudal gradient of self-similarity values in healthy controls, and an opposite gradient for multifractality (descending values, rostral to caudal). Schizophrenia patients had similar, although attenuated, gradients of self-similarity and multifractality values. Statistical tests showed that patients had higher values of self-similarity than controls in fronto-temporal regions, indicative of more regularity and memory in the signal. In contrast, patients had less multifractality than controls in the parietal and occipital regions, indicative of less diverse singularities and reduced variability in the signal. In addition, supervised machine-learning, based on logistic regression, successfully discriminated the two groups using measures of self-similarity and multifractality as features. Our results provide new insights into the baseline cognitive functioning of schizophrenia patients by identifying key alterations of criticality properties in their resting-state brain data.

## 1. Introduction

The global prevalence of schizophrenia is reported to be close to 21 million individuals (Charlson et al., 2018). The symptoms and poor prognosis of those affected can deeply impact their daily functioning, and weigh on those close to them. Unfortunately, progress in therapeutic development is slow in the field of psychiatry due to the extreme complexity of the brain, the heterogeneity of patients’ symptoms and difficulties in translational research. More knowledge is needed to better understand what alterations occur in the neural activity of patients. Among the missing pieces, further characterization of the resting neural dynamics of schizophrenia, and their relationship to patients’ symptoms, is needed. Alterations in the rhythmic (oscillatory) neural activity of schizophrenia patients have been widely reported in the neuroimaging literature (reviews: Uhlhaas and Singer, 2010; Maran et al., 2016; Alamian et al., 2017). In addition, an emerging body of research has reported changes in the arrhythmic properties of brain dynamics in schizophrenia (Breakspear, 2006; Fernández et al., 2013). A powerful concept that has so far remained under-exploited and poorly understood in neuropsychiatry is criticality.

### 1.1 What is criticality?

The dynamics of many complex systems, such as the human brain, appear to reside around the critical point of a phase transition (Beggs and Plenz, 2003; Stam and De Bruin, 2004; Fraiman and Chialvo, 2012; Palva and Palva, 2018). At this point of criticality, these systems are in a wavering state, at the cusp of a new phase, between the states of order and disorder (Beggs and Timme, 2012; Cocchi et al., 2017; Souza França et al., 2018). The brain requires such a balance of regularity (i.e., structure) on the one hand, to maintain coherent behaviour, and flexibility (i.e., local variability) on the other hand, to adapt to ongoing changes in the environment (R. Chialvo, 2004; Beggs and Timme, 2012). Indeed, critical brain dynamics have been shown to be optimal for fast switching between metastable brain states, for maximizing information transfer and information storage within neural networks (Socolar and Kauffman, 2003; Haldeman and Beggs, 2005), and for optimizing phase synchrony (Yang et al., 2012). Importantly, it is within a critical state that neural communication can span the greatest distance and achieve maximal correlational length (Fraiman and Chialvo, 2012). Thus, the brain’s state of criticality is thought to affect the functional properties of oscillations, local synchronization and signal processing (Palva and Palva, 2018). Changes to this state, due to psychiatric illness for instance, can alter certain properties of this balance (e.g., in terms of strength and number of synaptic connections) (Beggs and Timme, 2012). Some of the tuning parameters of criticality appear to be embedded in the balance between neural excitation and inhibition (e.g., through NMDA receptors (Mazzoni et al., 2007; Shew et al., 2009; Hobbs et al., 2010; Poil et al., 2012)), in neural network connection strengths, and synaptic plasticity (Rubinov et al., 2011; Beggs and Timme, 2012).

### 1.2 Measures of criticality

#### 1.2.1 Self-similarity and multifractality

Within the framework of criticality, local and large-scale fluctuations arise from excitatory post-synaptic potentials (EPSPs) and modulate brain states by facilitating or suppressing neuronal firing (Palva and Palva, 2018), with long-range spatial spread (He et al., 2010; Zilber, 2014). Systems in this state are characterized by power-law distributions, fractal geometry and fast metastable state transitions (Plenz and Chialvo, 2009; Cocchi et al., 2017; Chialvo, 2018; Palva and Palva, 2018). These features of a critical state are said to be scale-free or scale invariant. Power-law distributions of a given signal can be recognized as a linear slope in the log-log plot of the feature distribution, and they imply that the signal’s statistics and structural characteristics are preserved across spatiotemporal scales—in other words, that the signal has fractal properties (Beggs and Plenz, 2003; Chialvo, 2018). Fractal architectures describe objects that contain identical, or statistically-equivalent, repetitive patterns at different magnifying scales (Mandelbrot, 1983, 1985; Van Orden et al., 2012; Fetterhoff et al., 2015).

Scale invariant dynamics of systems at criticality (i.e., power-law distributions and fractal architecture) have often been described using a 1/f^β^ power law fitted to Fourier-based spectral estimations. On the other hand, self-similarity is a well-accepted model for scale-free dynamics and is richer than the sole measure of β, as it captures fractional Gaussian noise and fractional Brownian motion. Self-similarity can be measured by the Hurst exponent, *H*. In the brain, *H* is thought to index how well neural activity is temporally structured (via its autocorrelation). The smoother the signal, the higher the value of *H* (Zilber, 2014). However, self-similarity alone does not fully account for scale-free dynamics or criticality, since it can only capture additive processes (La Rocca et al., 2018). Combining self-similarity with multifractality improves on this framework to better capture criticality in a system. Multifractality can account for the remaining non-additive, non-Gaussian processes. The multifractality parameter, *M*, quantifies the diversity of *H’*s (singularities) and the overarching geometry of spatiotemporal fluctuations (Leonarduzzi et al., 2016; La Rocca et al., 2018). Generally, fractals are evaluated using the topological dimension, *D*, which describes the complexity and structure of an object by measuring the change in detail based on the change in scale (Di Ieva, 2016). In multifractal analysis, the local regularity of a signal is quantified using the *Hölder exponent, D(h)* (Jaffard et al., 2016), allowing a more realistic characterization of phenomena that are too complex to be explained solely by Euclidian models. In sum, the brain’s degree of criticality is defined by its scale-free dynamics, which are best quantified by combining measures of self-similarity and multifractality.

#### 1.2.2 Common measures of criticality

Numerous metrics have been developed to measure the scale-free properties that define criticality, such as Detrended Fluctuation Analysis (DFA) applied to oscillatory envelopes (Linkenkaer-Hansen et al., 2001a; Hardstone et al., 2012) and neuronal avalanche detection (Beggs and Plenz, 2003). Non-linear dynamics, and specifically multifractal analysis, has been used to address questions of self-similarity and multifractality. Multifractal analysis can characterize both the amount of global self-similarity in a system and the amount of local fluctuations (i.e., number of singularities) (Zilber et al., 2012). This approach allows for more in-depth interpretations of the electrophysiological data compared to more conventional analytical approaches. A number of mathematical frameworks have tapped into this, such as the Multifractal Detrended Fluctuation Analysis (MFDFA; (Kantelhardt et al., 2002; Ihlen, 2012)) and the Wavelet Leaders-based Multifractal Analysis (WLMA; (Wendt, 2007; Wendt and Abry, 2007; Serrano and Figliola, 2009)). For reviews of scale-free and multifractal analytical approaches, see (Lopes and Betrouni, 2009; Zilber, 2014).

#### 1.2.3 Application to psychiatry

A scoping review of alterations of brain criticality changes in clinical populations was recently discussed in Zimmern (2020). An insightful illustration of reported changes to the state of criticality across multiple neurological and psychiatric disorders, from the perspective of self-similarity, are illustrated in Figure 6 of that article (Zimmern, 2020). The application of criticality models to psychiatry, and in particular to the study of schizophrenia (SZ), is well in line with leading theories for this pathology, which are centered around dysconnectivity and altered information processing and transfer (Weinberger et al., 1992; Friston and Frith, 1995; Fernández et al., 2013). So far, most of the empirical evidence for dysconnectivity theory in SZ has come from functional magnetic resonance imaging studies, which highlight several important alterations in anatomical and functional connectivity that exist in SZ patients, as well as from electroencephalography (EEG) and magnetoencephalography (MEG) connectivity studies (review: (Alamian et al., 2017)). However, we still lack a complete, in-depth understanding of the brain alterations inherent to this pathology in the temporal domain.

In terms of scale-free analyses in psychiatry, power spectral densities (PSD) of resting-state fMRI scans have shown SZ patients to have reduced complexity and disrupted scale invariant dynamics compared to controls in the precuneus, inferior frontal gyrus and temporal gyrus, and these changes correlated with their symptoms (Lee et al., 2020). Electrophysiological studies have found altered dimensional complexity and increased variability in SZ patients’ signal (Koukkou et al., 1993). A number of studies have applied different versions of multifractal analysis on electrophysiological (Slezin et al., 2007; Racz et al., 2020) or white-matter MRI data in SZ (Takahashi et al., 2009). One of these used the multifractal analysis on resting-state EEG data, and found increased long-range autocorrelation and multifractality in patients compared to controls (Racz et al., 2020).

In addition, two insightful reviews have examined how non-linear methods could improve our understanding of SZ (Breakspear, 2006; Fernández et al., 2013). They highlighted conflicting results among studies reporting on complexity changes in SZ, which they proposed were attributable to participants’ symptomatic state, the method of imaging or medication. Complexity as measured by Lempel–Ziv complexity (LZC) or correlation dimension (D2) was typically found to be increased in SZ in studies that recruited younger, first-episode patients who were drug-naïve and symptomatic, while studies reporting SZ-related reductions in complexity tended to recruit older, chronic, patients who were on medication and hence less symptomatic (Lee et al., 2008; Fernández et al., 2013). Although these measures have been widely applied to neuroscientific data, they each come with caveats that affect their precision or generalizability. Moreover, these reviews highlight the importance of controlling for factors such as age and medication when studying complex pathologies, such as SZ.

### 1.3 Goals of the study

The brain is functionally optimal when in a state of criticality—in other words, when neural activity can spread equally well at long and short distances in time and space and information is processed and stored efficiently (Shew et al., 2009)—and multifractality analysis is among the most reliable indicators of criticality. Meanwhile, leading neural theories of SZ emphasize a pathological spatiotemporal dysconnectivity. It follows that the potential insights to be gained from multifractal analysis in SZ is extensive.

Therefore, the aim of the present study is to examine how criticality is altered in the neural activity of chronic SZ patients using the high temporal resolution of resting-state MEG data, the spatial resolution of structural MRI and wavelet-based estimations of multifractality and self-similarity. We hypothesize that, compared to healthy controls, chronic SZ patients will exhibit reduced self-similarity in the prefrontal cortex and increased self-similarity in sensory brain regions, along with reduced multifractality overall. We also predict significant correlations between measures of criticality and patients’ clinical symptom scores.

## 2. Materials and methods

### 2.1 Participants

Participant data collection was conducted at the Cardiff University Brain Research Imaging Centre in Wales, U.K., and the data analyses were conducted at the University of Montreal, QC, Canada. Ethical approval was obtained for the data collection according to the guidelines of the United Kingdom National Health Service ethics board, and the Cardiff University School of Psychology ethics board (EC.12.07.03.3164). Ethical approval was also obtained for these analyses from the research committee of the University of Montréal (CERAS-2018-19-069-D).

Behavioural and neuroimaging data from 25 chronic SZ patients (average age = 44.96 ± 8.55, 8 females) and 25 healthy controls (average age = 44.04 ± 9.20, 8 females) were included in this study. Healthy controls had no history of psychiatric or neurological disorders. The collected demographic information from all participants included: age, gender, depression score on the Beck Depression Inventory – II (BDI-II (Beck et al., 1996)), and mania score on the Altman Self-Rating Mania Scale (ASRM (Altman et al., 1997)). For the SZ patient group, additional information was collected: scores on the Scale of the Assessment of Positive Symptoms (SAPS) and the Scale of the Assessment of Negative Symptoms (SANS) (Kay et al., 1987), and information on antipsychotic doses standardized using olanzapine equivalents (Gardner et al., 2010). All of these data were anonymized, such that no identifiable information of participants was associated with their data nor with data from subsequent analyses. Patients were overall fairly asymptomatic on the testing day. No statistically significant group differences were observed across these demographic and clinical metrics, except for BDI-II scores, where SZ patients had on average mild depression (14.83 ± 9.11), compared to controls (4.50 ± 4.67). Additional details on participant information (i.e., recruitment procedure, exclusions, inclusions, and sample size calculation) can be found in Alamian et al. (2020) (Alamian et al., 2020).

### 2.2 MEG experimental design

The brain imaging data used for this study comes from five-minutes of resting-state MEG recorded during an eyes-closed condition, with a 275-channel CTF machine. Reference electrodes were placed on each participant to account for cardiac, ocular and other potential artefacts (Messaritaki et al., 2017). The MEG signal was initially recorded at a sampling frequency of 1200Hz. To conduct the MEG source reconstruction analysis, individual anatomical-T1 MRI scans were also recorded for each participant.

### 2.3 Data preprocessing and MEG source reconstruction

NeuroPycon (Meunier et al., 2019), an open-source python pipeline, was used for the preprocessing and source-reconstruction analyses. First, the continuous raw data was down-sampled from 1200Hz to 600 Hz, and band-pass filtered between 0.1-150 Hz. Next, independent component analysis (ICA) was used to remove artefacts from the MEG signal using MNE-python (Hyvarinen, 1999; Gramfort et al., 2013).

Since it has been reported that the values of the Hurst exponent, *H*, are unusually low in sensor-space, and tend to increase when moving from sensor to source space (based on simulations and real data: (Blythe et al., 2014)), source-reconstruction steps were taken to present cortical-level results in multifractal analysis. To generate individual anatomical source-spaces, the anatomical T1-MRI information of each subject was segmented with FreeSurfer (Fischl, 2012). However, given that this process would produce different source-space dimensions for each participant, individual source spaces were morphed and projected onto a standardized space from FreeSurfer (*fsaverage*) (Greve et al., 2013). The resulting source-space comprised 8196 nodes on the cortical surface, where dipoles were 5mm apart. The single layer model boundary element method implemented in MNE-python was used to compute the lead field matrix (Gramfort et al., 2013). Weighted Minimum Norm Estimate (Dale and Sereno, 1993; Hämäläinen and Ilmoniemi, 1994; Hincapié et al., 2016), implemented in the MNE-python package (Hyvarinen, 1999; Gramfort et al., 2013), was used to compute the inverse solution with a Tikhonov regularization parameter of *lambda* = 1.0 (Hincapié et al., 2016). Dipoles of the source-space were constrained to have an orientation perpendicular to the cortical surface. Thus, for this study, 8196 time series were extracted at the cortical level.

### 2.4 Characterization of criticality through self-similarity and multifractality

#### 2.4.1 Measuring self-similarity and multifractality

The singularity spectrum is a concise way to summarize information about scale-free dynamics. It allows the plotting of the Hölder exponents (*h*) about local variability in a time series, against the Fractional (Hausdorff) Dimensions, *D(h)*, as can be seen in Figure 1.

**Figure 1:**
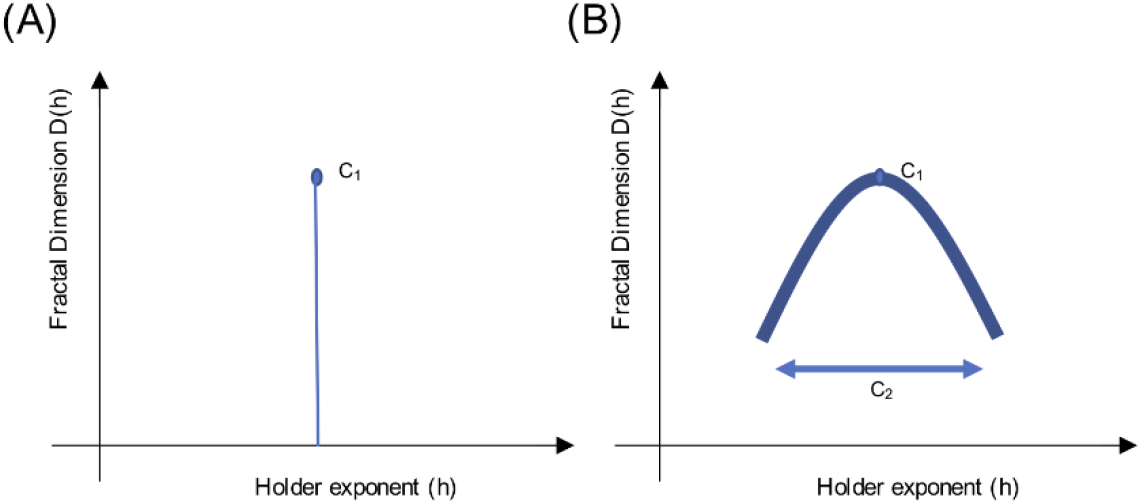
Sketch of a singularity spectrum. These sketches illustrate the multifractal scaling function, which depicts a singularity spectrum. Local variability in the signal is represented by Hölder exponents, h, on the x-axis, while the amount of singularities is represented by the Fractal Dimension, D(h), on the y-axis. The apex of the curve reveals the most common h exponent, while the width of the curve reveals the multifractal spectrum. Using log-cumulants from the WLBMF (described in 2.4.2) to describe the singularity spectrum, C1 informs on the apex, while C2 informs on the width of the function. Panel (A) shows a monofractal function, where C1=H and C2 = 0, and panel (B) shows a multifractal function, where the concavity shows the distribution of h singularities.

Multifractal analysis builds on measures of self-similarity (e.g., slope of the PSD, DFA) to provide information about local fluctuations (singularities) in time. The multifractality spectrum and the scaling function ζ(q) (in terms of statistical moments *q*) are related, and can be described using the Legendre transformation:

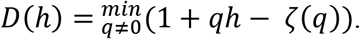

When a signal is monofractal, this becomes a linear function, where ζ(q) = qH, as it would only have a single singularity (one unique property, Figure1a). Here, the self-similarity parameter would be equal to *H*, the Hurst exponent (Wendt et al., 2007). When a signal is multifractal, the function ζ(q) has a curvature, as in Figure 1b, which shows the global spectrum of singularities. The Hölder exponent (*h*) with the largest Fractal dimension, *D*, (apex of the curve) is said to be the most common singularity in the time-series. The width of the curve can be described with the multifractality parameter, *M* (Wendt et al., 2007).

In this study, to meaningfully estimate self-similarity and multifractality, we used the Wavelet p-Leader and Bootstrap based MultiFractal analysis (WLBMF). This approach builds on the Wavelet leaders-based multifractal analysis (WLMA) method that has been thoroughly described elsewhere (Wendt, 2007; Wendt et al., 2007; Serrano and Figliola, 2009; Ciuciu et al., 2012; Fetterhoff et al., 2015). Briefly, this WLMA method of estimating the singularity spectrum was shown to be efficient in untangling the scaling properties of neuronal signal, and more robust than other algorithms in addressing non-stationarity issues (Wendt, 2008). The curved shape of the scaling function ζ(q) can be written in its polynomial expansion around its maximum to allow the evaluation of *C*_*p*_, log-cumulants:

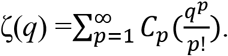

The singularity spectrum can be thus derived from the series-expansion of *C*_*p*_. The first two log-cumulants are the most informative, with C1, the first log-cumulant, reflecting self-similarity (and the location of the maximum of *D(h)*, similar to *H*). Its values approximate those of the *H*; values above 0.5 indicate positive correlation (signal has memory), values below 0.5 indicate negative correlation, and a value of 0.5 indicates lack of correlation (random signal). Meanwhile, C2, the second log-cumulant, reflects multifractality (and the width of the singularity spectrum, like *M*) (Wendt, 2007; Wendt et al., 2009; Zilber, 2014; Diallo and Mendy, 2019). Given the concavity of the scaling function, C2 is always negative, and when C2 equals 0, it is said to indicate monofractality.

Hölder exponents cannot take on negative values. Thus, most multifractal analyses are constrained to scaling functions that have only positive local regularities, implying that there is a continuous temporal positive correlation in the signal (i.e., locally bound everywhere in the function). However, this is not true of all brain signals, which can present with discontinuities in the signal and can thus take on negative regularities. Thus, p-leaders have been proposed as a way to circumvent this limitation (full description in : Jaffard et al., 2016).

#### 2.4.2 Defining parameters of log-cumulants

One method to detect criticality in the brain is through the Wavelet p-Leader and Bootstrap based MultiFractal (WLBMF) analysis and, more specifically, through the evaluation of log-cumulants (Wendt and Abry, 2007; Wendt et al., 2007). This MATLAB-implemented technique uses the discrete wavelet domain for the analysis of self-similarity and multifractality in signals. In order to compute C1 and C2 in our study, we first plotted the PSD of each participant group (SZ patients, controls) in log-log space and identified the portion of the PSD function exhibiting a log-linear relationship. In our data, the log-linear portion of the PSD belonged to j1=7 and j2=10, which correspond to 3.5 Hz and 0.4 Hz, respectively, as deduced by the following equation: 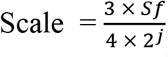, where Sf represents the sampling frequency, j1 and j2 represent the start and end points of the log-linear portion, respectively, and the scale represents the frequency bin to which it corresponds. This frequency range is similar to those of other researchers who have used the same multifractal analysis (Zilber, 2014). The PSD was calculated at the overall cortical level and also at the ROI level, using the Destrieux Atlas (Destrieux et al., 2010), to ensure that the linear part of the spectrum remained the same across brain regions. For the purposes of this study, we used second order statistics in the evaluation of the log-cumulants (i.e., p-leader of p=2), corresponding to long-range temporal correlations computed with DFA (Leonarduzzi et al., 2016). For the ROI-based investigations, the C1 and C2 log-cumulants were first computed for each node (n=8196 sources) in cortical source-space, and then averaged across ROIs (n=148 ROIs based on the Destrieux atlas, (Destrieux et al., 2010)).

### 2.5 Statistics and machine-learning analyses

#### 2.5.1 Conventional statistics and correlation analyses

Group statistical analyses were conducted between SZ patients and matched-controls to test for group-level differences in C1, C2, and demographic and clinical data. This was done at the ROI and source levels. To do so, we used non-parametric statistical tests (two-tailed, unpaired, pseudo t-tests), corrected with maximum statistics using permutations (n=1000, p<0.001) (Nichols and Holmes, 2001; Pantazis et al., 2005).

Moreover, Pearson correlations with False Discovery Rate (FDR) correction (Genovese et al., 2002) were used to explore the relationship between cortex-level C1/C2 values and scores on the SANS, SAPS and medication-dosage, in patients. FDR correction (Benjamini-Hochberg) was applied to each p-value (computed for each of the 8196 nodes) to account for the multiple comparisons in order to achieve a significance threshold of p<0.05, corrected.

#### 2.5.2 Machine learning analyses

MEG signal classification was conducted using a logistic regression model and a stratified 10-fold cross-validation scheme to evaluate the discriminative power of the log-cumulants C1 and C2 in classifying SZ patients and controls. First, at each of the 8196 nodes, the feature vector (either C1 or C2 values), computed for each participant, was split into 10 folds, while maintaining a balance between the two classes (SZ and controls). Next, the classifier was trained on the data from nine of the ten folds and tested on the remaining fold (test set). The classification performance was assessed using the decoding accuracy (DA) on the test set (i.e., percentage of correctly classified participants across the total number of participants in the test set). This operation was repeated iteratively until all the folds were used as test sets. The mean DA was used as the classification performance metric. In order to infer the statistical significance of the obtained DAs, permutation tests were applied to derive a statistical threshold as described in (Combrisson and Jerbi, 2015). This method consists of generating a null-distribution of DAs obtained by running multiple instances of the classification (n=1000), and randomly shuffling class labels each time. Maximum statistics were applied in order to control for multiple comparisons across all the nodes (Nichols and Holmes, 2001; Pantazis et al., 2005). Visbrain was used for all the ROI and cortical-level visualizations (Combrisson et al., 2019).

## 3. Results

### 3.1 Alterations in self-similarity and multifractality

The group averages of C1 and C2 values for SZ patients and healthy controls can be seen in Figure 2. Across both participant groups, a clear gradient in C1 values was observed, where self-similarity values increase gradually from the frontal lobe to the occipital lobe. Interestingly, a similar gradient, but in the opposite direction, is observed in terms of C2 values in both groups, with C2 values gradually increasing from the occipital lobe to the frontal lobe. Moreover, the magnitude of this gradient appears less pronounced in patients than in controls.

**Figure 2:**
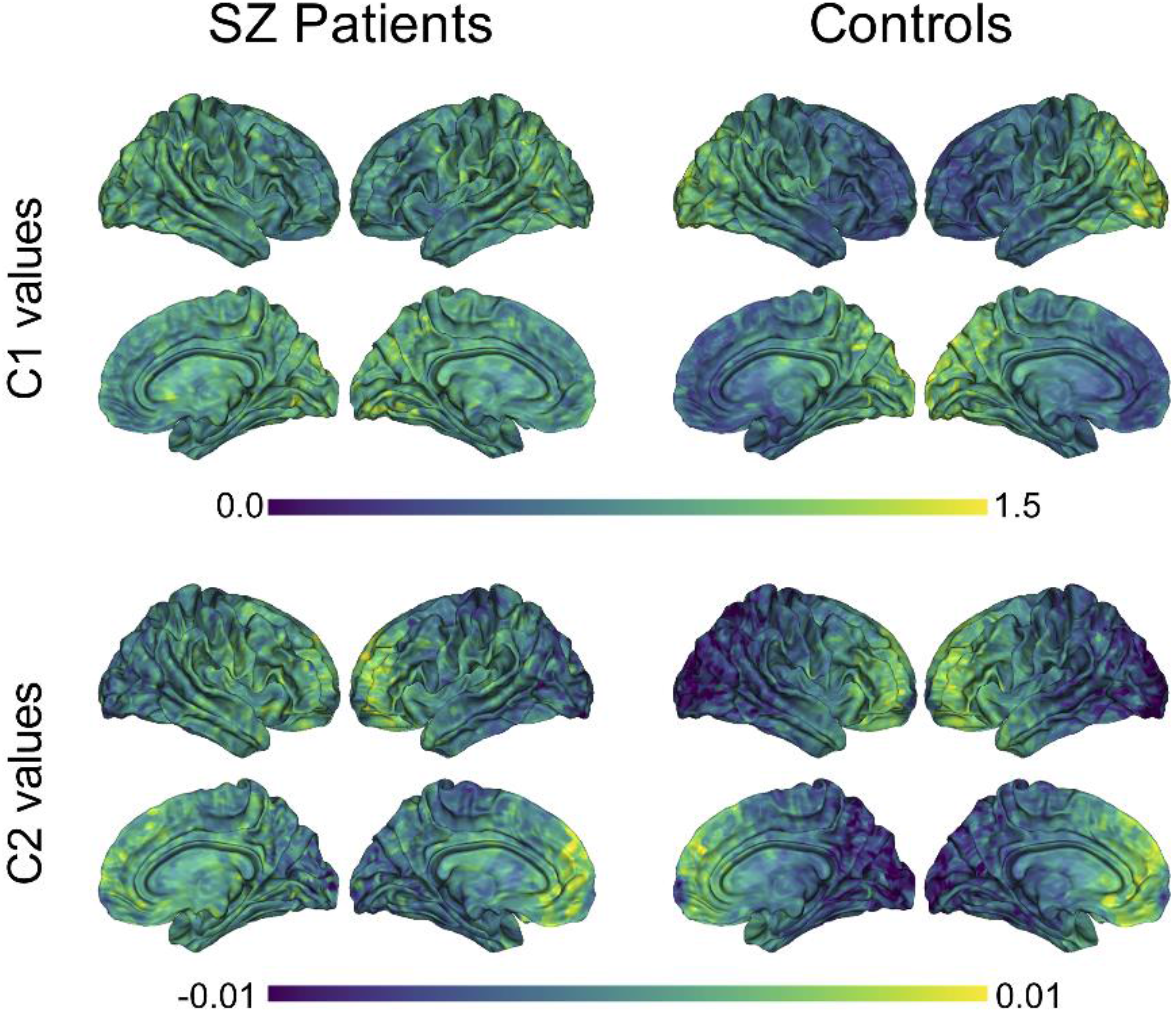
Group averages of C1 and C2 values in SZ patients and controls. Averaged C1 and C2 values were computed for each of the 8196 nodes, within each group. P-leader p=2 was used. SZ= schizophrenia

Conventional unpaired t-tests between the two subject groups did not yield any statistically significant differences in terms of C1 or C2 values (p<0.05, two-tailed t-test). Figure 3a shows t-values for the direction and magnitude of group differences for C1 and C2 values, where positive (red) t-values indicate brain areas where SZ patients have smaller C1 or C2 values compared to controls, and negative (blue) t-values indicate brain areas where patients have larger C1 or C2 values compared to controls.

**Figure 3:**
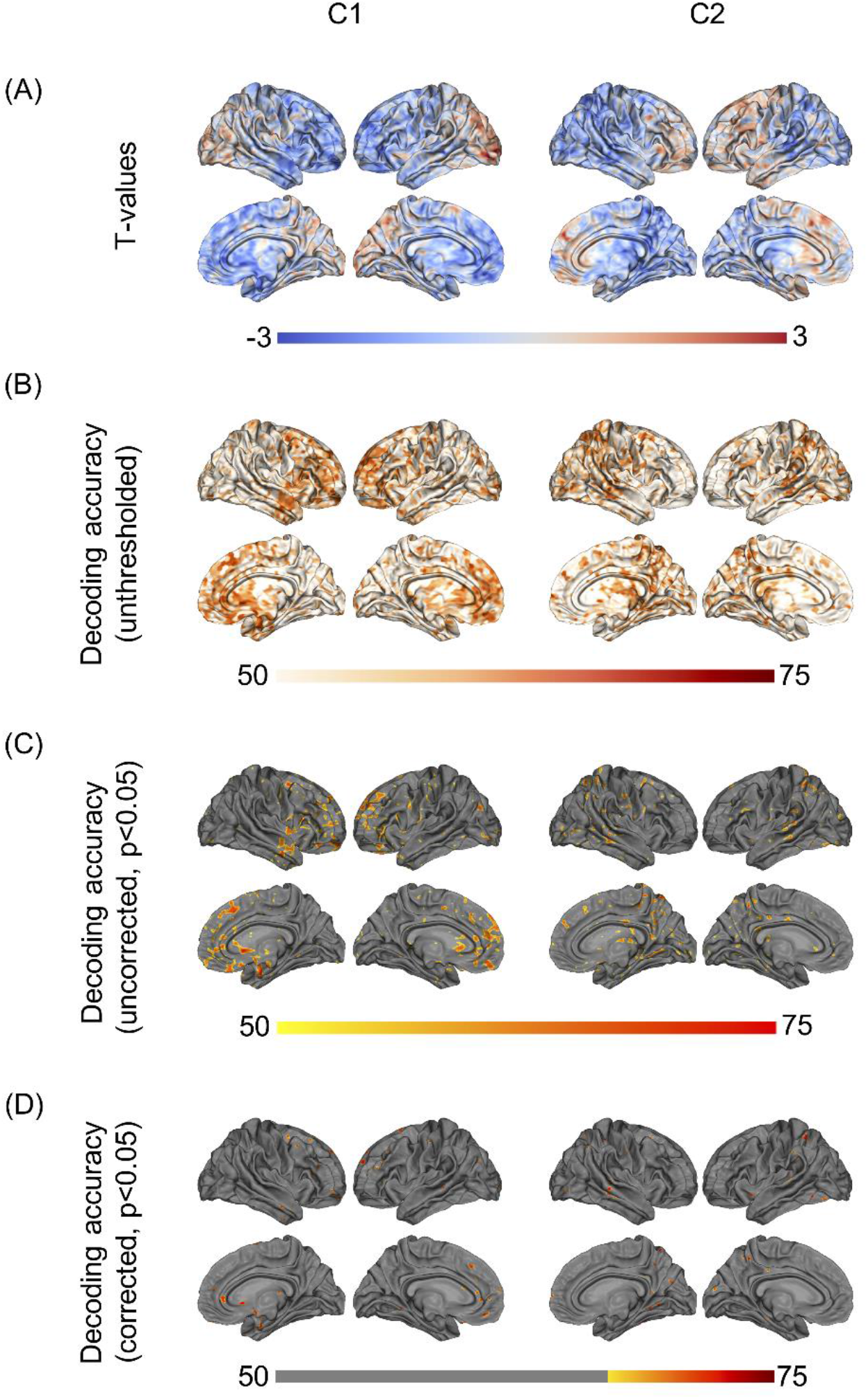
Group differences and machine-learning results. (A) shows t-values from the unpaired t-tests (non-significant), showing (controls – patients). (B) shows unthresholded DA values based on logistic regression, using C1/C2 as a single feature. (C) shows the same DA values, thresholded at p<0.05. (D) shows the DA values corrected for multiple comparisons using maximum statistics (p<0.05), thresholded at the chance level of 70%. P-leader p=2 was used. DA= decoding accuracy.

By contrast, when using a machine-learning approach to test for out-of-sample generalization in the same data, we found that C1 and C2 in multiple brain regions led to statistically significant classification of the two subject groups, with up to 77% decoding accuracy (Figure 3d, max statistics correction, p<0.05). More specifically, using source-space C1 values as a decoding feature led to statistically significant discrimination of SZ and controls in the subcallosal gyrus, middle fontal gyrus and anterior part of the cingulate gyrus, bilaterally. The left superior frontal gyrus, the left inferior frontal gyrus and sulci, and the right orbital, straight and frontomarginal gyri were also significant. The maximum decoding occurred in the left superior frontal gyrus (77%, compared to the chance level of 70%). Meanwhile, using source-space C2 values as a decoding feature led to statistically significant classification of SZ patients and controls in the superior parietal lobule, precuneus and posterior-ventral part of the cingulate gyrus in the right hemisphere. The left post-central gyrus, and superior temporal gyrus and occipital gyrus, bilaterally, were also significant. The maximum decoding accuracy took place in the right temporal gyrus (76%, compared to the chance level of 70%). Figures 3b-d show the unthresholded DA values for C1 and C2, as well as the uncorrected results at p<0.05, and the corrected classification results at p<0.05, with multiple comparisons correction using max statistics.

Figure 4 shows the classification results based on C1 and C2 values computed at the ROI-level (p<0.05, corrected for multiple comparisons). The ROIs involved in the significant discrimination of patients and controls were the left straight gyrus, the triangular part of the inferior frontal gyrus and the medial transverse frontopolar gyrus and sulcus for C1, and the superior occipital gyrus, the right cuneus and the left angular gyrus for C2. To illustrate how the classifier was able to successfully separate SZ patients from healthy controls, individual C1 and C2 values were computed and averaged across all brain sites that had shown significant decoding at the source-level. These values are presented in a scatter plot in Figure 5. The distribution of the individual C1 and C2 values (averaged over all sources with significant decoding accuracy) shows that C1 values are higher in patients than in controls (i.e., a trend towards more self-similarity) and C2 values also shift upwards in patients (i.e., a trend towards less multifractality).

**Figure 4:**
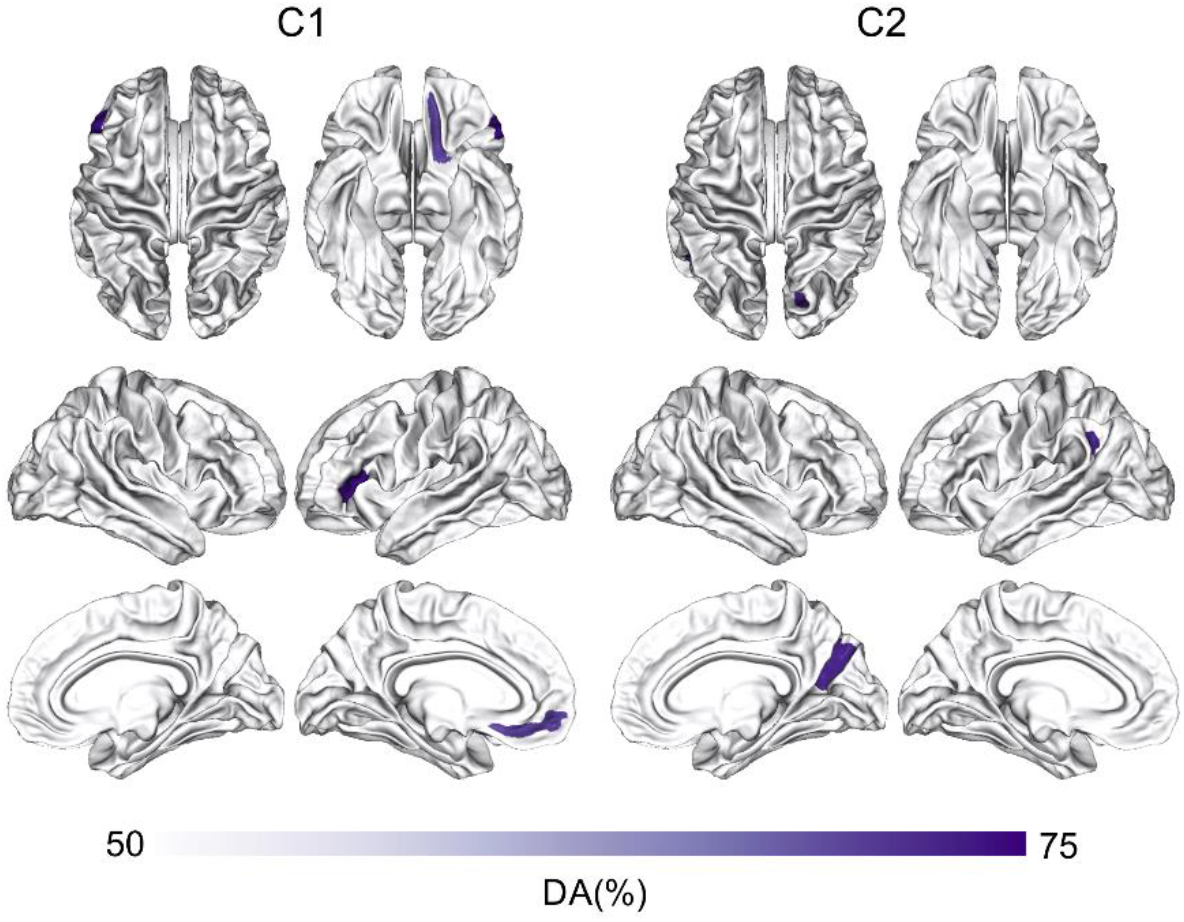
ROI-based classification of SZ and controls using C1 and C2. Machine-learning classification of SZ patients and healthy controls using logistic regression and the features of C1 or C2 at the ROI-level. The ROI analysis was based on the Destrieux Atlas, p<0.05, corrected for multiple comparisons. DA=decoding accuracy, SZ=schizophrenia.

**Figure 5:**
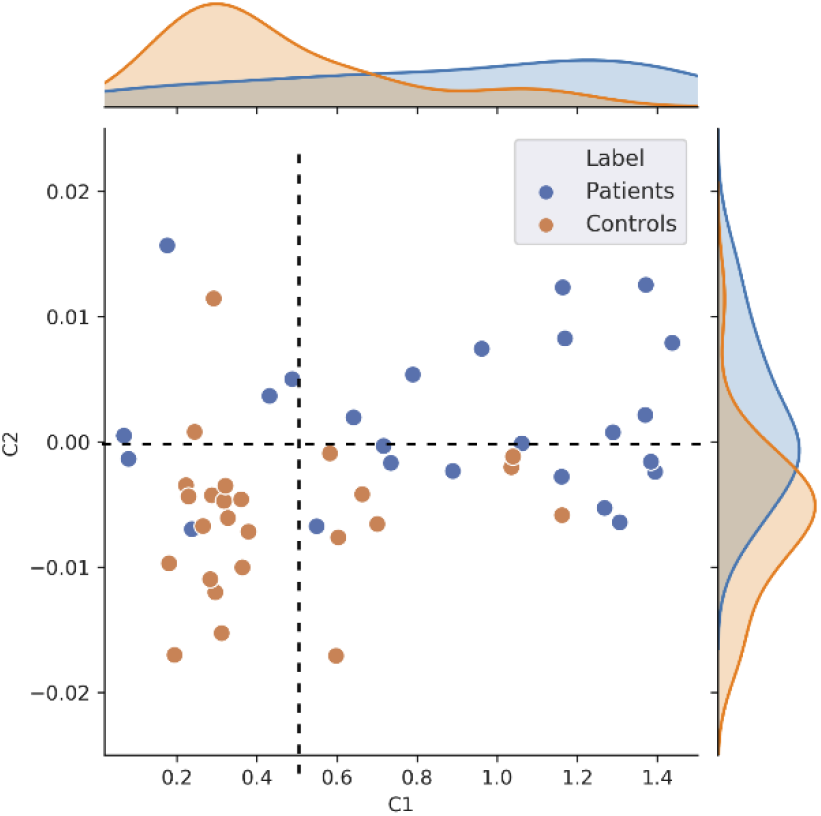
Scatter plot visualization of individual C1 and C2 values. This figure shows individual C1 and C2 values, averaged across all the nodes that showed statistically significant patient vs controls decoding (n=50). This scatter plot illustrates that patients exhibit overall higher self-similarity (higher C1) and less multifractality (higher, less negative, C2).

**Figure 6:**
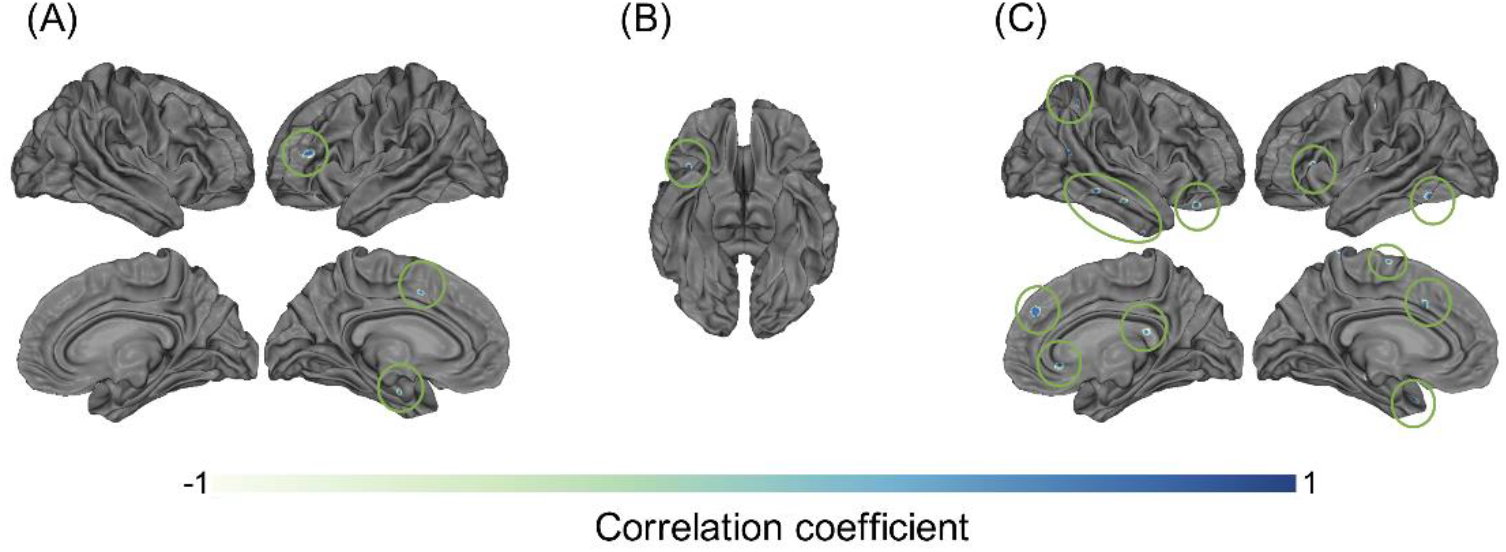
Correlational results between C1 and C2 values and patients’ clinical information. Pearson correlation results between patients’ (A) C1 values and negative symptom scores on the SANS (p<0.05), (B) C2 values and positive symptom scores on the SAPS (p<0.05), and (C) C1 values and medication dosages (olanzapine equivalent in mg), p<0.05.

It is noteworthy that this scatter plot reveals the presence of positive C2 values in the dataset, primarily in patients. Although mathematically ill-defined, the observation of positive C2 is not unprecedented. Positive C2 values in some individuals can be attributed to numerical instabilities (and might be statistically undistinguishable from 0) or to the fact that the data in these participants cannot be modeled using the multifractal formalism. The safest interpretation for the positive C2 values observed in Figure 5 (primarily in patients), is that data in these individuals were neither multifractal (C2 < 0) nor monofractal (C2 = 0). Given that this specific type of multifractal analysis has never been conducted on clinical data before, we explored how the results would change when using a p-leader of p=4 (as opposed to the p=2 we have used up to now). This analysis found fewer participants to have positive C2 values compared to p=2, and generally allowed for a better modeling of multifractality in the resting neuromagnetic signal of participants. Figures of C1/C2 group averages and classification patterns based on p=4 can be found in the Supplementary material 1. In summary, we observed a similar albeit stronger decoding of patients and controls based on C2 values in p=4 than p=2. Interestingly, C1 values were smaller (Supp. Fig. 1), and the strong frontal lobe classification results based on C1 values at p=2 diminished at p=4 (Supp. Fig. 2 c-d). Taken together, the results of C1 estimation (self-similarity) were more reliable in our data when using a p-leader of p=2, while C2 estimation (multifractality) provided more robust results with p=4. Most importantly, the trends in terms of increasing C1 and C2 values in patients compared to controls was present irrespective of the choice of p.

### 3.2 Correlations with clinical scores/information

The investigation of potential correlations between C1/C2 and clinical information resulted in a number of interesting results. Specifically, the correlations between C1 values and patients’ SANS scores (maximum r=0.78, p<0.05) in the left inferior frontal gyrus and sulcus (Figure 6a), and between C2 values and patients’ SAPS scores (maximum r=0.78, p<0.05) in the circular sulcus of the insula (Figure 6b) were statistically significant. In addition, the relationship between C1 and medication dosage yielded a statistically significant positive correlation (maximum r=0.79, p<0.05, after correcting across all nodes). Figures 6c and 7c illustrate that patients with higher medication dosage exhibited higher C1 values. This was especially significant in the superior frontal gyri, the right middle temporal gyrus, left mid-anterior cingulate gyrus and left inferior temporal sulcus (see Figure 6c). The positive correlations in these analyses are shown in the scatter plots in Figures 7a-c. These plots depict the relationship between individually-averaged C1 and C2 values (based on the significant nodes), and patients’ symptom severity and medication dosages. To further clarify the C1 x SANS correlational results, a Pearson correlation was conducted between SANS scores and medication dosage, revealing a low-to-moderate correlation coefficient. The r2 of the regression model suggested that this relationship explained 27-40% of the data, meaning that the correlation of C1 x SANS was only partially mediated by medication.

**Figure 7:**
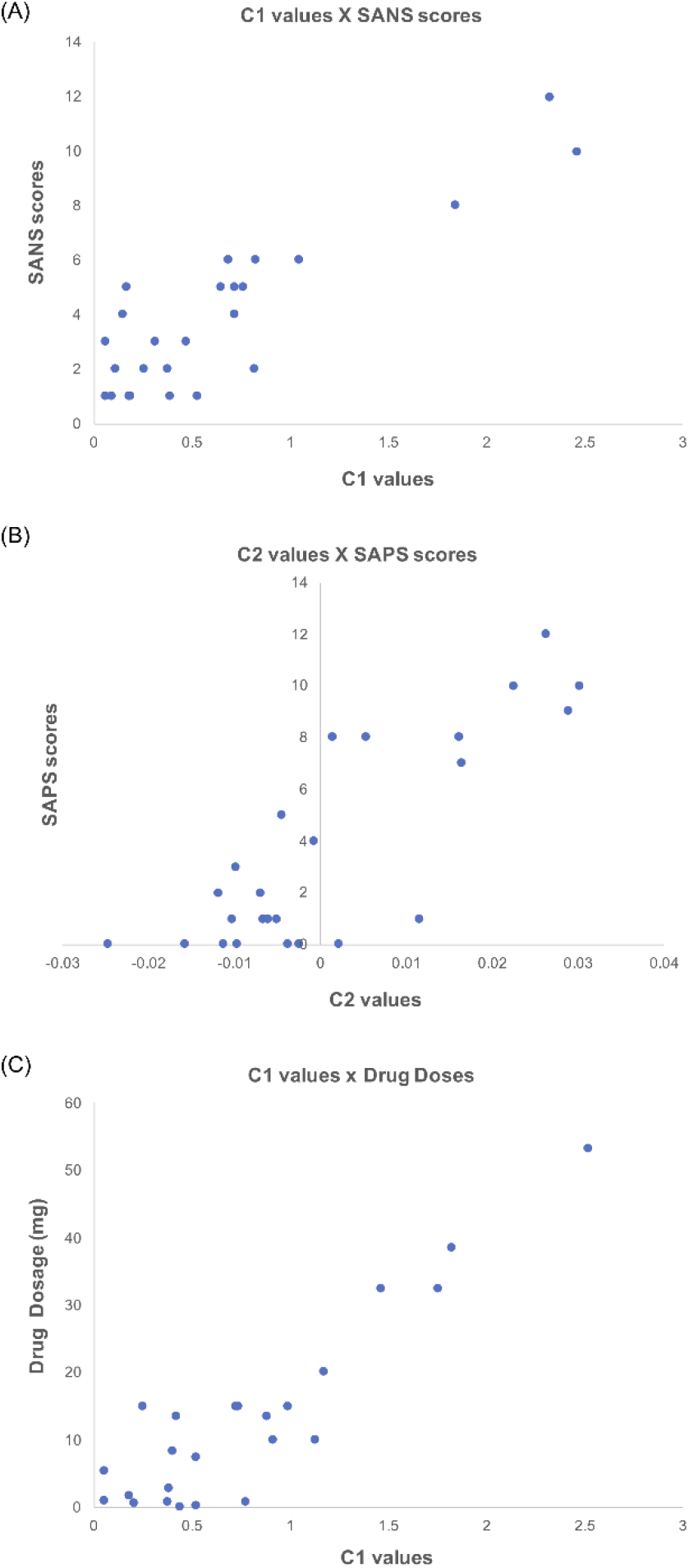
Scatter plots showing the positive correlations between C1/C2 values and clinical information. These scatter plots depict correlations (p<0.05) between individually averaged C1 and C2 values and subjects’ clinical information. The averaging of C1 and C2 values was over significant nodes. (A) Shows the correlation between C1 values and patients’ SANS scores, (B) shows the correlation between C2 values and SAPS scores, and (C) shows the correlation between C1 values and patients medication dosage (olanzapine equivalent in mg).

## 4. Discussion

The central goal of this study was to examine and characterize criticality features in the baseline neural dynamics of schizophrenia. To do so, we evaluated the first two log-cumulants of the Wavelet p-Leader and Bootstrap based MultiFractal (WLBMF) analysis on the resting-state neuromagnetic signals of chronic SZ patients and healthy controls. This allowed us to determine the values of C1 (reflective of self-similarity) and C2 (reflective of multifractality) on the linear, scale-free portion of participants’ arrhythmic MEG signal in source-space. In brief, our findings partially supported our initial hypotheses about self-similarity and multifractality changes in SZ, whilst also revealing unexpected alterations in criticality.

Specifically, the findings of this study show that there are clear opposite gradients in the values of C1 and C2, along the rostro-caudal axis. A progression from low to high values of C1 were observed from anterior to posterior poles (i.e., frontal to occipital lobes), while C2 values showed the reverse progression. For both of these metrics, the gradient was less clear in SZ patients than in healthy controls. The t-values of the unpaired t-tests showed that patients had higher C1 values in the fronto-temporal area, and lower C1 values in the parieto-occipital areas compared to controls. In contrast, patients appeared to have higher C2 values in the temporal, parietal and occipital areas than controls. Conventional t-test statistics failed to reach significance after multiple comparisons correction. However, a machine-learning approach based on logistic regression yielded statistically significant decoding (up to 77%) of patients and controls in a number of brain regions. Indeed, SZ patients and controls were categorized using C1 values in the anterior part of the cingulate gyrus (ACC), the left inferior gyrus, and the mid and superior frontal gyri, among other brain regions. Meanwhile, using C2 as a feature, we were able to statistically significantly classify patients and controls in the right temporal gyrus, precuneus, and occipital gyrus, among other brain regions.

In terms of the first log-cumulant, patients had a range of C1 values of [0.07,1.44] in significant regions. In controls, this range was of [0.18,1.16]. Typically, C1 (and thus *H*) values would be expected to be between 0 and 1 (where 0 < C1 < 0.5 implies negatively autocorrelated signal, C1 = 0.5 implies uncorrelated signal, and 0.5 < C1 < 1 implies positively autocorrelated signal), although values above 1 have been observed within the theory of *generalized processes and tempered distributions* (Samoradnitsky and Taqqu, 1994). In terms of the second log-cumulant, patients had a range of C2 values of [-0.01,0.015] in significant brain regions, while controls had a range of [-0.02, 0.011]. These values fall within the same ranges reported by previous researchers (e.g., (Zilber, 2014). As a reminder, higher C1 values are indicative of more self-similarity and memory in the signal, while lower (more negative) C2 values are indicative of more complexity in the form of multifractality. From our results, we infer that SZ patients exhibited more self-similar neural dynamics than healthy controls, and thus more regularity in the frontal and temporal brain areas. In addition, patients had fewer singularities (less diverse *h)* in the parietal and occipital brain regions, compared to healthy controls whose neural signals were more multifractal.

Further investigation of this analysis revealed that a subportion of participants (predominantly patients) had some positive C2 values. Theoretically, only [C2 < 0] (multifractal signals) or [C2 = 0] (monofractal signals) are expected. Observing positive C2 values implies that the multifractal formalism could not properly model the neuromagnetic data recorded in these patients. So, what does this tell us about the success of the classifier in using C2 to distinguish between patients and controls? The simplest explanation is that individuals with more negative C2 (stronger multifractal properties) were identified as healthy, whereas individuals with C2 values closer to zero (monofractal), or even higher than zero (neither multifractal no monfractal), were classified as patients. As a side note, we found that using an alternate p-leader of p=4 improved C2 values, and the classifier reaffirmed the diminished multifractality characteristics of patients’ resting neuromagnetic signal. Taken together, we observe clear rostro-caudal gradients of ascending self-similarity and multifractality across both participant groups, albeit more clearly in controls. The reduced multifractality and increased self-similarity might reflect a certain rigidity in the temporal dynamics of SZ patients’ neural activity.

Our findings are consistent with recent publications that have characterized complexity in SZ in the same regions in which we observed alteration in the log-cumulants C1 and C2 (i.e., precuneus, inferior frontal gyrus and temporal gyrus, (e.g., Lee et al., 2020)). Interestingly, a recent resting-state MEG-based study of SZ patients by La Rocca et al. (2018) also found a gradient in C1 values along the longitudinal axis; however, in contrast to our own finding of an ascending anterior-posterior gradient, they instead found an opposite, descending anterior-posterior gradient (La Rocca et al., 2018). In addition, La Rocca et al. (2018) compared how criticality features changed during a perceptual task. They reported that in healthy individuals, global self-similarity decreased, while focal multifractality increased when switching from rest to task. Moreover, the changes in multifractality correlated with brain regions implicated in the task. This finding could suggest that the metric of C2 has a functional role in cognitive processes (La Rocca et al., 2018). Of note, there are some methodological differences between our studies, such as the choice of scale (j1 and j2) for the linear portion of the PSD. Differences could also be due to age differences. Indeed, the authors reported the mean age of their participants to be 22 years old, while our group’s mean age was of 44 years old. In the complexity literature, it has been often reported that the properties of scale-free dynamics change with age (e.g., Fernández et al., 2011; Churchill et al., 2016), and so it is possible that there is a reversal of the self-similarity gradient with age. More work is needed to elucidate this.

Positive correlations were observed between the metrics of self-similarity and multifractality and patients’ clinical information. In particular, we observed an increase in C1 values in patients with increasing severity of scores on the negative symptoms scale (SANS) in the inferior frontal gyrus, as well as with patient’s medication dosage, the latter of which was especially strong (r=0.79). The left frontal gyrus plays an important role in cognitive functioning (Swick et al., 2008) and language (Klaus and Hartwigsen, 2019). At the structural level, cortical thinning has been observed in the inferior frontal gyrus in SZ patients compared to healthy controls, which correlated with cognitive dysfunction (Kuperberg et al., 2003; Oertel-Knöchel et al., 2013). Correlation between inferior frontal gyrus volume and negative symptoms in SZ patients have been previously observed, but not in their non-affected siblings (Harms et al., 2010). At the functional level, higher cluster coefficients have been observed in the left inferior frontal compared to bipolar patients or controls (Kim et al., 2020), as well as weaker connectivity within the language network (Jeong et al., 2009). In addition to the reported structural alterations in this language processing center, the reduction in the temporal flexibility and enhanced regularity in the signal might explain why patients’ have poorer speech understanding, such as difficulty detecting metaphors, sarcasm or jokes (Rossetti et al., 2018). A correlational trend was also observed between multifractality and patients’ scores on the positive symptom scale (SAPS) in the circular sulcus of the insula. In past studies, negative correlations have been observed between reduced grey matter volume of the insula and SZ patients’ positive symptoms (Wylie and Tregellas, 2010; Cascella et al., 2011). It is interesting to note that self-similarity and multifractality were oppositely (and perhaps complementarily) correlated with symptom severity scores.

Taking into account the correlational findings, it is not surprising that, in our dataset of chronic and medicated patients, antipsychotic medication dosage was related to symptom severity, which itself was related to scale-free neural properties. Psychiatrists typically increase pharmaceutical dosage, gradually and as needed, to help manage symptoms. Sometimes, certain drug combinations that help manage positive symptoms (hallucinations, delusions) can worsen negative symptoms (Schooler, 1994; Goff et al., 1996). Evidence from other studies (Koukkou et al., 1993; Saito et al., 1998; Raghavendra et al., 2009) suggests that drug-naïve and first-episode patients may display a different pattern of criticality, thus the generalizability of our results is limited to other medicated, chronic SZ patients.

Another parallel can be drawn between this study’s results and findings from DFA analyses. The log-cumulants (C1 and C2) derived from WLBMF analysis using a p-leader of p=2, as was used in the present study, are similar to scaling exponents obtained using DFA (Leonarduzzi et al., 2016), in that they both reflect temporal autocorrelations. In one of our recent publications, we computed DFA exponents on oscillatory envelopes in this same dataset of SZ patients and healthy controls (Alamian et al., 2020). The scale used for the computation of the log-cumulants (j1,2: 0.4-3.5 Hz) overlaps with the delta oscillatory band (0.5-3.5Hz). Comparing delta DFA exponents and C1 between the studies reveals a good agreement: DFA exponents were reduced in patients compared to controls in the occipital and parietal lobes and increased values in the prefrontal and temporal lobes, similar to the C1 topology. The overlap was remarkably good considering that DFA was computed on band-limited rhythmic brain signal, while the log-cumulants of the singularity spectrum were computed on the arrhythmic raw brain signal. This comparison shows that while DFA is an adequate measure of the self-similarity aspect of criticality, it does not however provide any information on the multifractality of a signal, as does the second log-cumulant, C2. In this respect, they capture different properties of the neural signal, and should be treated as such. Several studies have examined the alterations of DFA across a number of psychiatric and neurological disorders. They found that a drop in DFA exponents occurs in SZ as well as in Alzheimer’s and Parkinson’s disease, whereas other conditions, such as depression, insomnia and epilepsy are typically associated with increases in DFA exponents (Zimmern, 2020). These findings reveal that reduced temporal autocorrelations observed in SZ are not disease-specific, but capture alterations that might be common to multiple psychiatric or neurological conditions. This again highlights the need for more elaborate measures of brain criticality, such as through the WLBMF analysis carried out in the present manuscript.

Criticality in the brain likely informs on the spatiotemporal organization and functioning of neural networks at the micro- and macroscopic levels (Hesse and Gross, 2014; Cocchi et al., 2017). While the origins of criticality are still debated, many agree that scale-free neural fluctuations are the signature of a brain in a state of criticality. A right balance of scale invariant properties (self-similarity, multifractality) is thought to be needed to adapt and respond to ever changing environments (Linkenkaer-Hansen et al., 2001b; Plenz and Chialvo, 2009; Beggs and Timme, 2012; Palva et al., 2013; Shew and Plenz, 2013). Consequently, we propose that a change in this equilibrium could disrupt optimal brain functioning. When self-similarity is strong in a signal, as in the brain signals of our SZ cohort, the signal’s temporal autocorrelation decays slowly, such that signal memory lasts a long time. While still the subject of debate, it has been proposed that this enhanced temporal persistence (or redundancy) may make the brain less efficient in information processing (Zilber et al., 2013). Lower levels of self-similarity in signals, as in those of our healthy controls, are thought to reflect enhanced neural excitability and more efficient processing (He, 2011, 2014; Zilber et al., 2013). Interpretations of multifractality are still unclear, but it appears that a richer repertoire of singularities (multifractality > monofractality) suggests more variability and flexibility in the neural signal (Beggs and Timme, 2012), and thus in behaviour. In our dataset, patients exhibited reduced multifractality in certain areas, thus suggesting a decrease in complexity and flexibility in their resting neuromagnetic signal. The observed alterations in these criticality metrics in SZ could explain the long, sustained nature of patients’ positive symptoms (delusions, hallucinations) and their difficulty in breaking away from them.

## Conclusion

The overarching scale invariance of brain activity is thought to be a useful indicator of its organization across both temporal and anatomical scales (Werner, 2007; Zilber, 2014). Indeed, many have suggested that biological systems optimally process, adapt to and communicate information over long neural distances when in a state of criticality. This critical state involves a balance between regularity (structure) and flexibility (variability, local fluctuations). Disruption of this equilibrium may reduce the efficiency with which the system responds to changes in the environment. In this study, we applied WLBMF analysis to resting MEG signals and observed clear deviations in both the self-similarity and multifractality of these signals in chronic SZ patients compared to healthy controls. These changes in the state of criticality of patients lend further support to the theory of dysconnectivity in SZ from the perspective of temporal dynamics, as it characterizes a different way in which information interruption occurs in patients. This study also demonstrated that alterations in neural criticality can be used to accurately differentiate between chronic SZ patients and controls. We expect that these findings will fuel the search for strong biomarkers in SZ, borrowing a new, largely uncharted path.

## Acknowledgements

We would like to thank Dr. Herwig Wendt for helpful discussions and Jordan O’Byrne for helpful comments on the manuscript.

## Supplementary Material

**Supp. Fig.1.**
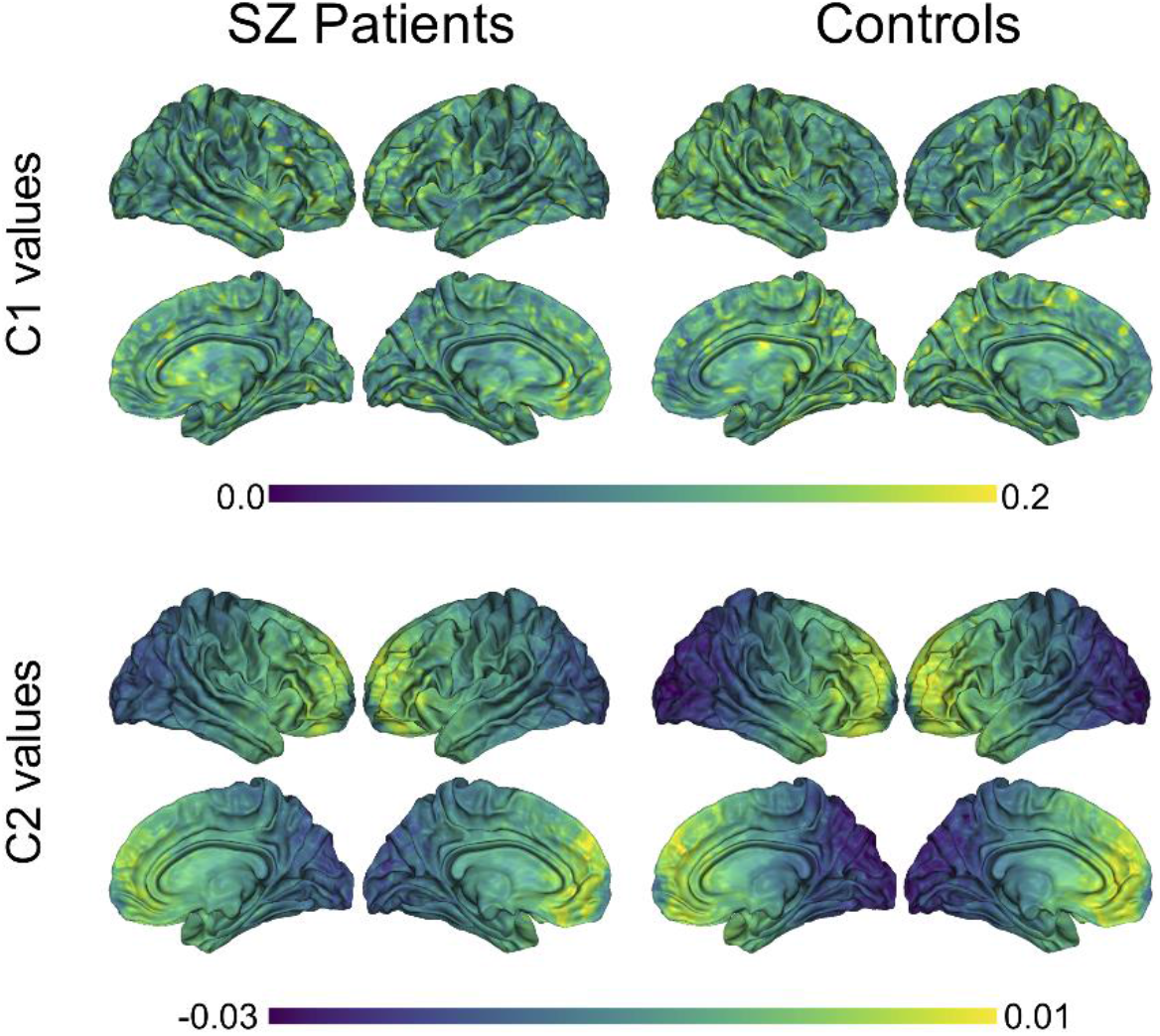
Group averages of C1 and C2 values in SZ patients and controls. Averaged C1 and C2 values were computed for each of the 8196 nodes, within each group. P-leader p=4 was used. SZ= schizophrenia

**Suppl. Fig.2.**
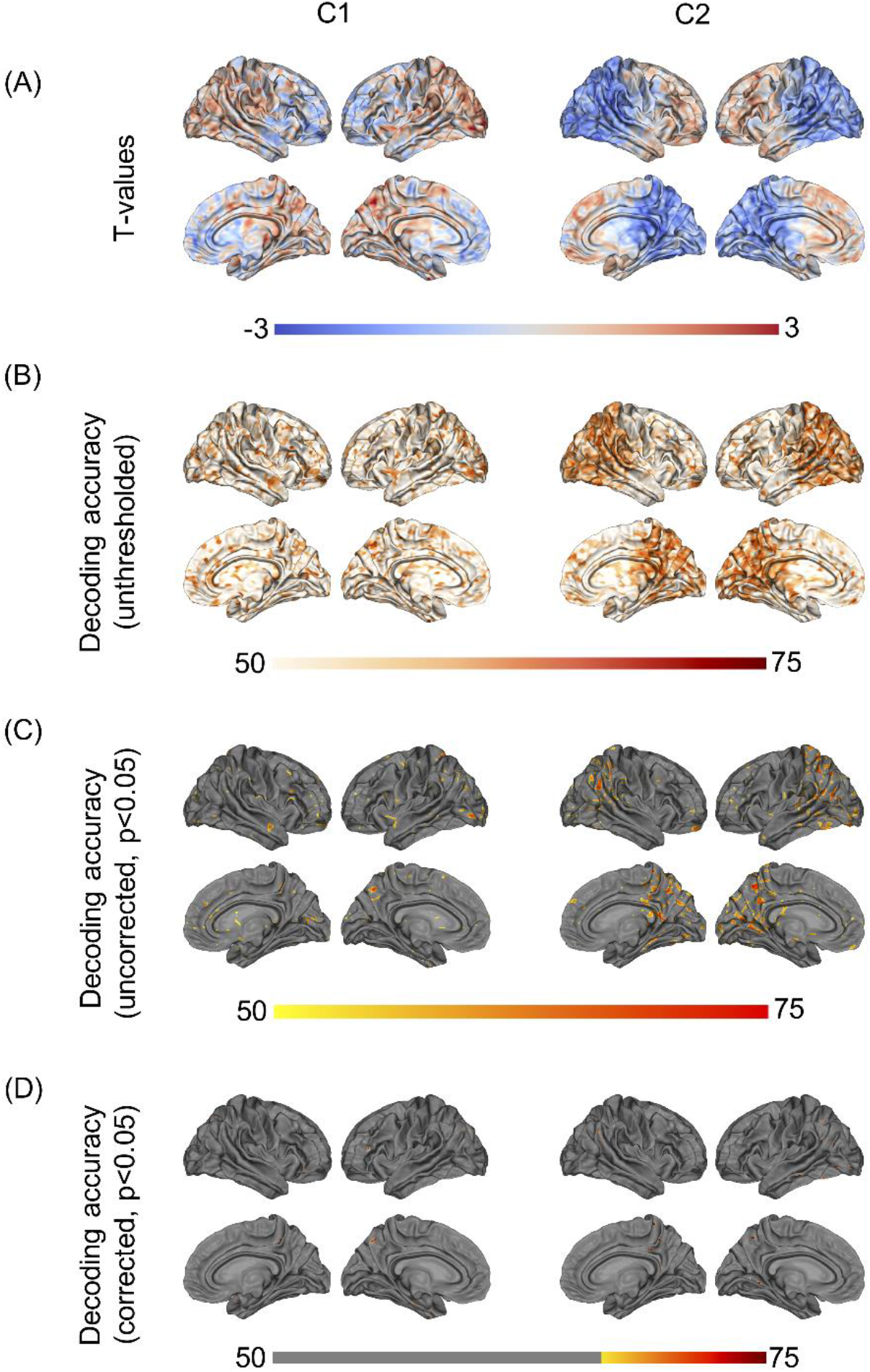
Group differences and machine-learning results. (A) shows t-values from the unpaired t-tests (non-significant), showing (controls – patients). (B) shows unthresholded DA values based on logistic regression, using C1/C2 as a single feature. (C) shows the same DA values, thresholded at p<0.05. (D) shows the DA values corrected for multiple comparisons using maximum statistics (p<0.05), thresholded at the chance level of 70%. P-leader p=4 was used. DA= decoding accuracy.

